# Correspondence between Signaling and Developmental Patterns by Competing Cells: A Computational Perspective

**DOI:** 10.1101/2023.05.22.541859

**Authors:** Zahra Eidi, Najme Khorasani, Mehdi Sadeghi

## Abstract

Arrangement of variant phenotypes in ordered spatial assemblies during division of stem cells is essential for the self-organization of cell tissues. The cellular patterns of phenotypes competing for space and resources against one another are mostly driven by secreted diffusible chemical signaling clues. This complex process is carried out within a chronological framework of interplaying intracellular and intercellular events. This includes receiving external stimulants-whether secreted by other individuals or provided by the environment-interpreting these environmental signals and incorporating the information to designate cell fate. An enhanced understanding of the building blocks of this framework would be of help to set the scene for promising regenerative therapies. In this study, by proposing a designative computational map, we show that there is a correspondence between signaling and developmental patterns that are produced by competing cells. That is, the model provides an appropriate prediction for the final structure of the differentiated cells in a competitive environment. Besides, given that the final state of the cellular organization is known, the corresponding regressive signaling patterns are partly predictable following the proposed map.

**Author Summary:** Multicellular organisms are made of repeated divisions of single cells and aggregation of their offspring together. However, the aggregated formations are not colony-like accumulations of piled-up cells. Instead, they are “emergent” spatiotemporal structures of developmentally differentiated cell types. The functionally integrated structures remain relatively constant throughout the life of the organisms, despite the death and production of new cells. The question is: How differentiated cells are capable of making variant patterns without any predefined templates? It is shown that with a variety of differentiated cell types, emergence of complex patterns is feasible through the interplay of intercellular interactions and intracellular decision-making switches. Such conceptual understanding has the potential to generate a multitude of novel and precisely controlled cellular behaviors.

## Introduction

The duality of variety and organization is among canonical concerns in biology. During the course of development, cells are subject to a diverse range of chemical stimuli, as underlying drivers of non-genetic variation, acting at multiple scales^1^. These stimuli play a crucial role in directing cell fate determination in stem cells at the individual cell level^2^. On the other hand, collective processes such as tissue homeostasis, wound healing, angiogenesis, and tumorigenesis are intimately linked with competing environmental chemical cues^3^. While, in principle, the genetic material of every single cell of an organism is the same, influenced by variant positional stimulants, they are capable of generating highly complex spatial patterns^4–7^. Understanding the mechanisms underlying the generation and maintenance of these ordered spatial assemblies could potentially aid in the development of novel strategies for controlling tissue organization and function in vitro and in vivo. The questions we would like to answer, or, more realistically, get any enlightenment about, are threefold; Firstly, In the presence of variant positional cues, how can spatially organized populations give rise to and maintain large-scale inhomogeneities starting from an initially roughly homogeneous mass of intermixed stem cell populations? Secondly, how do individual stem cells perceive and interpret their surrounding spatial information to make decisions about their developmental pathway in response to the local concentration of these stimulants? And finally, is it possible to infer information about the specific form of the signals that created them from the final structure of cell populations?

The basis of cellular pattern formation is mounted on the interaction of mediating nonlinear diffusive signaling components^8^. For the spontaneous construction of patterns during development, as proposed by Turing’s classic theory, the system requires two diffusive chemical compounds: an activator compound and an inhibitor one^9^. The latter locally undergoes an autocatalytic reaction to generate more of itself and also activates in some way the formation of the inhibitor compound. Meanwhile, the former inhibits the formation of more activator compounds. The key element for obtaining spatial patterns is that the activator component and the inhibitor one diffuse through the reaction medium at different rates. Thus, the effective ranges of their respective influences are different. Accordingly, if the inhibitor agent diffuses faster than the activator one, a stable pattern can emerge from a homogeneous background merely by amplification of small perturbations. The patterns, generally, take the form of spots (and reverse spots) or stripes based on the choice of model parameters^8^. The dynamic elaborates different possible pattern formation processes in a variety of developmental situations. The related examples span from the regeneration of hydras^10^ to animal coating patterns^11^. Wave phenomena also can generate patterns of spatiotemporal type^12,13^. Since the typical characteristic time of cell division is higher than that of a traveling wave, here, we exclude the formation of cellular patterns induced by spatiotemporal signaling patterns. Recently, Marcon *et al*. ^14^ proposed a new development in classical Turing models, indicating that the essential prerequisite of varied diffusion rates for mobile signaling molecules is not essential for pattern formation. Remarkably, specific networks are capable of creating patterns using signals without the constraint of relative diffusion rates.

Internal mechanisms are responsible for generating the right proportion of different types of specialized cells, distributing them into their right position, and maintaining the organized structure in the presence of intercellular chemical signaling agents^15^. Cells also sense and respond to mechanical stimuli and the physical properties of their environment via induced downstream genetic regulatory networks^16–18^. Several multi-stable regulatory networks play their role as the internal decision-makers of dividing cells^15^.

This study investigates the impact of various chemical signals on the mechanism by which multiple stem cells generate intricate tissue structures and tries to provide a deeper understanding of the mechanisms behind morphological variation. In reality, the formation of intermediate structures during embryo development or the formation of a tissue consisting of cells with different phenotypes and with organization in their spatial arrangement without a previous template is a complex problem, whose modeling using the simplest possible assumptions can lead to a better comprehension of the development process in multicellular organisms. We assume that there are two multipotent stem cells as resources of variation generation, each of which is potentially capable of constructing its own organized structure in the absence of the other. Here, we present a computational model for their internal mechanism in the presence of each other to form an organized population consisting of the whole descendants. We see that signaling messengers play a significant and irreplaceable role, as regulatory agents in communication between different species. Our results indicate that the association of variant environmental signaling messengers and intracellular decision-making switches grants a diverse range of cellular patterns. Furthermore, having the ultimate arrangement of cellular organization, one can approximately indicate the signaling patterns based on which the cellular patterns have been established, provided that the prior assumption of the pattern is given.

## Materials and Methods

Let us consider a plane which is partially covered by a population of two type of stem cells, *SC*_1_ and *SC*_2_. Here, to reduce the computational costs and without lose of generality, we assume that the stem cells directly divide into their corresponding differentiated cells and overlook the phase of all intermediate stages. We suppose that the stem cells are capable of both renewing themselves and dividing to their corresponding differentiated offspring. *SC*_1_ divides either to *A* and/or *B* and *SC*_2_ divides either to *C* and/or *D*. The amount of signaling agent that each individual cell receives affects on the likelihood of its division outcomes. Here, the main idea is that in the absence of cellular displacement, competition between existing chemical signals in the environment plays the principal role in pattern formation process at population level. To model the underlying mechanism, we need to answer the following questions:

- What type of signal the model does refer to?
- How can competing signals impact on an individual cell’s fate?
- What is the impact of signals on offspring at population level?

### Signals

Let us assume that the stem cells in a medium are exposed to special chemical information, we refer them as signals, that is captured and interpreted by them to develop spatial organization. There are various ways to provide spatial patterns in biology, among which positional information and reaction-diffusion dynamics are the most prominent ones^19^.

### Positional Information Dynamic

Generally, positional information dynamic is referred to the development of spatial cellular organization in the embryo differentiating at specific positions based on their response to gradient of environmental signals^3^. *E*.*g*., embryonic organizer centers, secrete morphogens that specify the emergence of germ layers and the establishment of the body’s axes during embryogenesis^20^. In the current study, by positional information we mean any external chemical cues whose procedure of setting-up is immaterial for us and we merely focus on their impact on regulation of internal switches. To illustrate the rivalrous relationship between different signals Fig.1 exemplifies the competition of two signal profiles of Gaussian type (the first column), a Gaussian profile and a sinusoidal one (the second and third columns) and two sinusoidal with different frequencies (the forth column).

**Figure 1.**
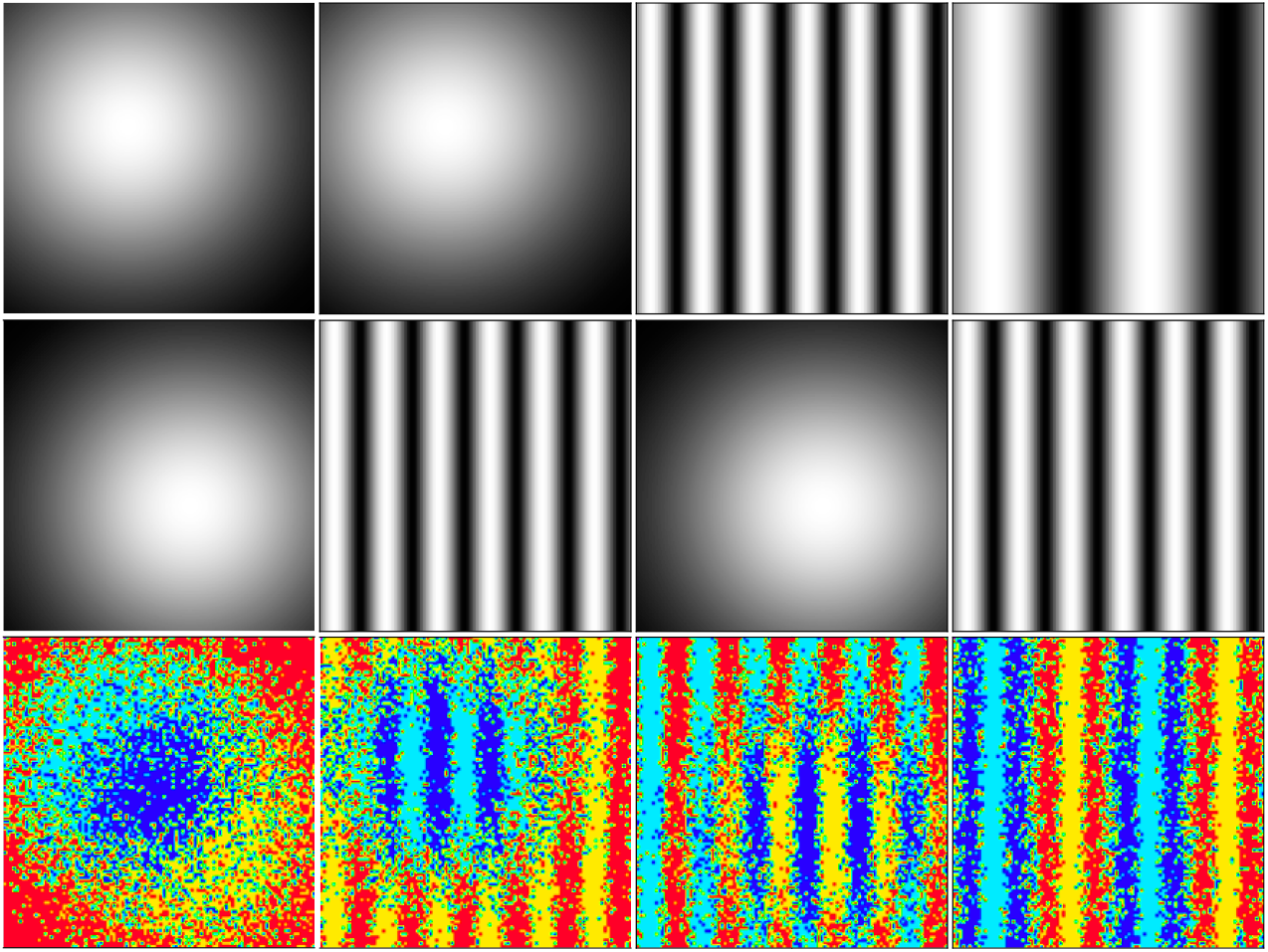
The resulted cellular pattern in an environment which is exposed to two independent signal patterns: A Gaussian profile with relation exp[(2*σ*2)^*−*1^((*x− x*^*∗*^)^2^ + (*y −y*^*∗*^)^2^)] and a sinusoidal one *sin*(*kx*). Here, the final result is determined based on the comparison of the signal concentrations at each point. The randomness involved in the patterns belongs to the areas where the concentration of positional signals are comparable. The first column: (Top) *x*^*∗*^ = 40, *y*^*∗*^ = 30 and *σ*= 2 (Middle) *x*^*∗*^ = 60, *y*^*∗*^ = 30 and *σ*= 2 (Bottom) Developed pattern in consequence of the combination of its upper-head signals.

The second column: (Top) *x*^*∗*^ = 40, *y*^*∗*^ = 30 and *σ*= 2 (Middle) *k* = 4.5 (Bottom) Developed pattern in consequence of the combination of its upper-head signals. The third column (Top) *k* = 4.5 (Middle) *x*^*∗*^ = 40, *y*^*∗*^ = 30 and *σ*= 2 (Bottom)Developed pattern in consequence of the combination of its upper-head signals. The forth column: (Top) *k* = 1.5 (Middle) *k* = 4.5 (Bottom) Developed pattern in consequence of the combination of its upper-head signals.

### Signaling through Reaction-Diffusion Dynamic

Suppose that there are two independent signaling agents in the medium, each of which is generated via its corresponding reaction-diffusion dynamic. Each reaction-diffusion system that we currently take advantage of it, involves two generic chemical species 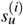 and 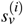, whose concentrations at a given point in space is specified by the same variables 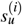 and 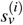 respectively.

We assume that the concentration field of 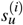 in the medium, *i∈*1, 2, defines the dynamic profile of one of the independent signaling agents. Moreover, 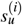 and 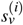 are deemed to spread over the environment with 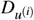 and 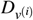 respectively. The governing equations to the propagation of the signals read as:

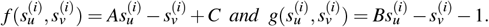

where, by rescaling the space variable, the diffusion coefficient of 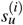 and 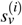 are set to 1 and *d*, respectively. Here, *d* is equal to 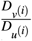 Thus, assuming that *d ≥* 1, the diffusivity of 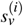 is larger than that of 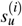. Besides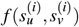and 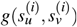 are reaction kinetics of the system represented with the following terms:

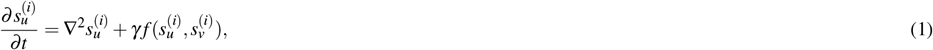

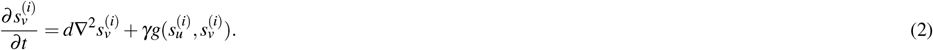

where, *A, B* and *C* are controlling parameters. The kinetics also constrains the variable 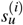 within a finite range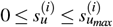 The parameter *γ* exhibits the relative strength of reaction kinematics. This dynamic with reflective boundary condition can produce steady state heterogeneous spatial patterns of chemical concentrations^21^. Diffusion process, with *d ≥* 1, in this context is considered as the main deriving process for the heterogeneity in the system. Moreover, 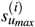 is considered as the controlling parameter upon which the behavior of spatial patterns differs, see Fig.2. In order to simulate the dynamic, we implement the Gillespie method^22^ which exhibits some degree of randomness in the simulation of chemical kinetics. This method generates a statistically possible solution of the master equations Eqs.1 and 2 for which the reaction rates are known. Defining the *propensity function* for every single reaction, including diffusion ones which are considered to be reducible to an analogous reaction, we have a measure to find out the time when the next chemical reaction takes place, and also which reaction is likely preferred by the system. The entire reactions of the system and their corresponding propensity functions are listed in the table.1. By updating the propensity functions at each step, one can track the changes in the corresponding species’ population vector which is induced by a single occurrence of a particular reaction. Repeating the algorithm simulates the whole behavior of the reaction-diffusion system, stochastically.

**Figure 2.**
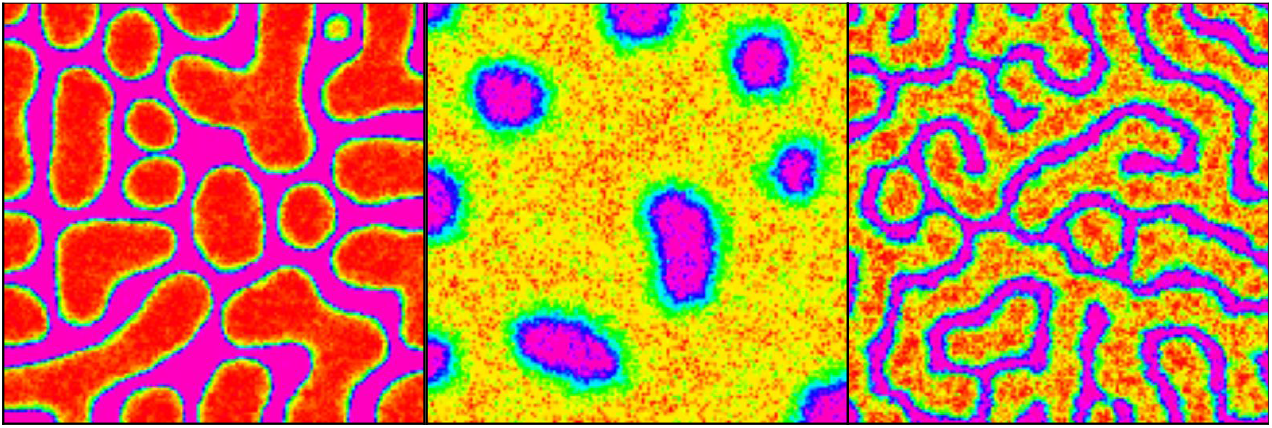
The whole possible patterns following the dynamic of Eqs.1 and 2 with parameters: *A* = 0.9, *B* = 1.2, *γ* = 10000, *C* = 0.2 and the size of lattice *h* = 0.01 (Left) Spot, *D*_*u*_ = 1 and *D*_*v*_ = 20 *D*_*u*_, (Middle) Reverse Spot, *D*_*u*_ = 25 and *D*_*v*_ = 500 *D*_*u*_ and (Right) Stripe, *D*_*u*_ = 1 and *D*_*v*_ = 20 *D*_*u*_.

**Table 1.**
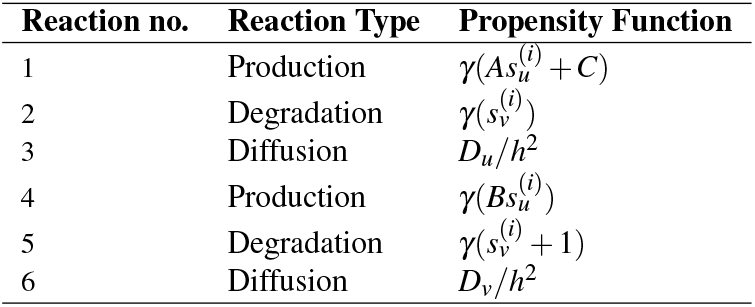
The involved reactions of the system which result in producing signaling patterns along with their corresponding propensity functions

| Reaction no. | Reaction Type | Propensity Function |
| --- | --- | --- |
| 1 | Production | $\gamma(As_u^{(i)} + C)$ |
| 2 | Degradation | $\gamma(s_v^{(i)})$ |
| 3 | Diffusion | $D_u/h^2$ |
| 4 | Production | $\gamma(Bs_u^{(i)})$ |
| 5 | Degradation | $\gamma(s_v^{(i)} + 1)$ |
| 6 | Diffusion | $D_v/h^2$ |

### Biased Internal Switch of Determinants

Specifying the environmental clues that can influence on the fate of stem cells brings us to the second question: How competing signals can impact on an individual cell’s fate?

Let us assume that the fate of each stem cell, *SC*_*i*_ (*i* = 1, 2), is controlled by a tri-stable regulatory switch, see Fig.4(a) :

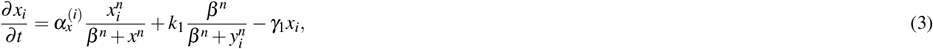

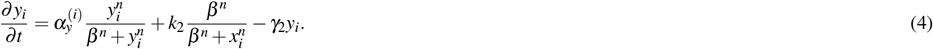

in which *x*_*i*_ and *y*_*i*_, *i ∈*1, 2, are cytoplasmic determinants of the stem cell *i*, whose their mutual interaction, Eq.3 and Eq.4, impels the cell to one of its probable fates. The dynamic includes mutual repression of *x*_*i*_ and *y*_*i*_ and their degradation effects as well as their self-activation in the form of Hill-function. In the above equations *n* is the Hill coefficient, *β* is synthesis rate of determinants, 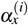 and 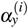 are self-activation rates, *k* = *k* are inhibition rates,and *γ* = *γ* are the degradation rate of *x* and *y*, respectively. This sort of dynamic for regulatory internal switches has vast precedent in the literature^23–25^. In the current study, we assume that 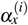 and 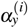 are not fixed parameters. Instead, influenced by competing environmental signals *s* and *s*, play a pivotal role in controlling the cell’s fate. The below relations governs behavior of 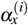 and 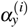

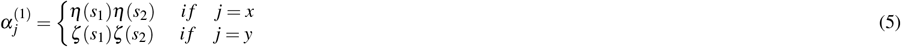

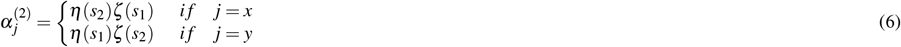

where 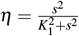 and 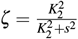 and *K*_2_ are fixed parameters.

### Patterns at Population Level

In this section, we present the position-dependent algorithm for an illustrative example depicted in Fig.3 by which the patterns of differentiated cells are developed at the population level. Let us assume that the system involves two different stem cells *SC*_1_ and *SC*_2_, each of which divides and develops into its corresponding differentiated cells according to the amount of signaling agents *s*_1_ and *s*_2_. *SC*_1_ is capable to differentiate into *A* and *B* phenotype whereas *SC*_2_ is competent to develop into *C* and *D* offspring, see Fig.4(b). *s*_1_ and *s*_2_ independently propagate on the substrate via reaction-diffusion dynamics of Eqs.1 and 2. The objective is to track the potential fate of stem cells at each location on a two-dimensional grid based on their exposure to two types of signals, *s*_1_ and *s*_2_. To achieve higher accuracy, the signal levels are scaled up to a range of 0 to 5. For each present species, a 6 *×*6 lookup table is created at the start of the simulation, where each element in the table represents the potential number of cells of the corresponding species after division, assuming that a specific combination of *s*_1_ and *s*_2_ signals (*s*_1_, *s*_2_) exists at the mother cell’s location.

To simulate the dynamic of the pattern formation through the division process provoked by the positional chemical information, we perform the following steps recurrently on a substrate with size *sz* = 100 on which *SC*_1_ and *SC*_2_ has been distributed randomly (Fig.3, panel of Initial Population Arrangement):

**Figure 3.**
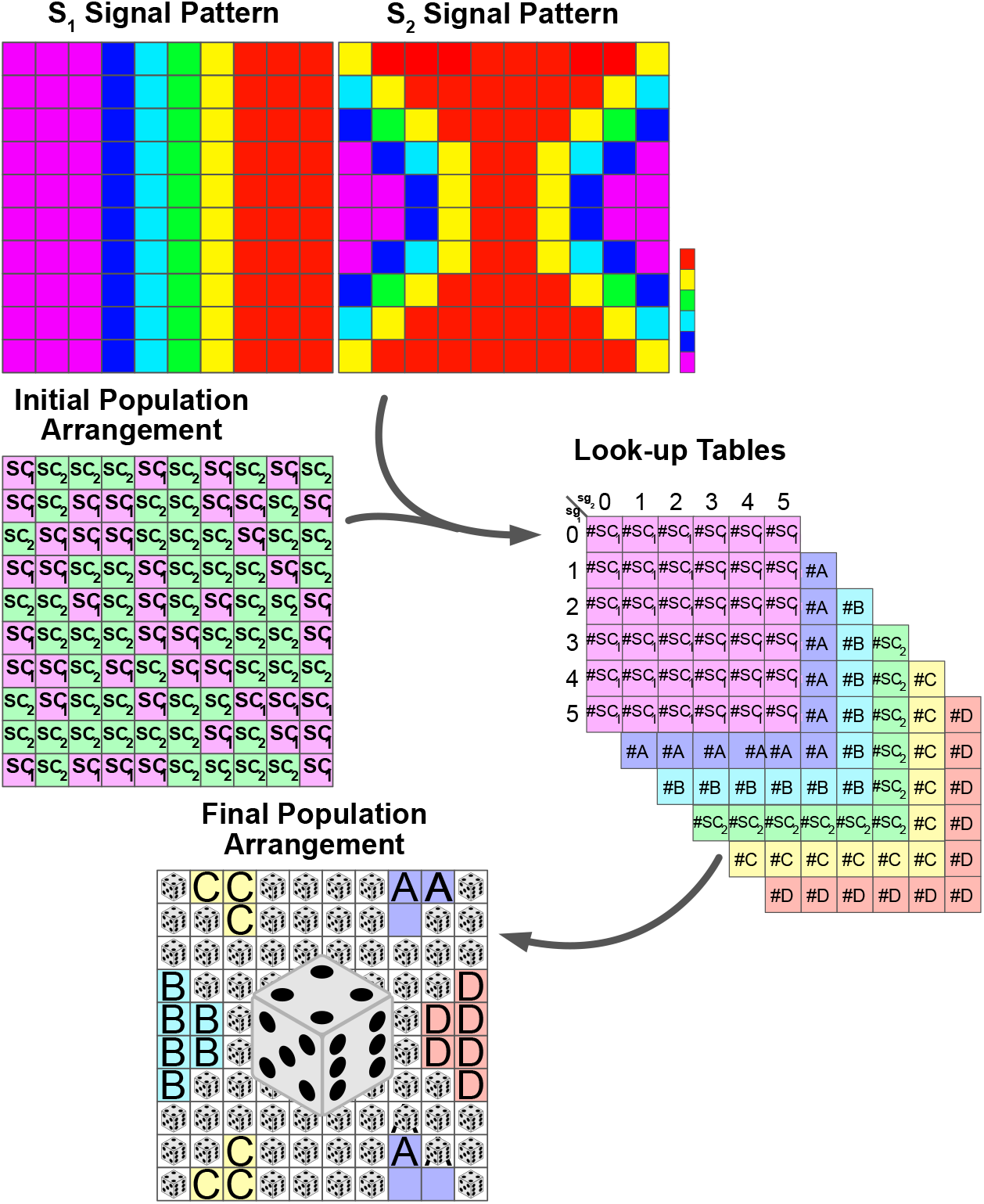
Schematic illustration of the chronological process of forming developmental patterns in accordance with the competing external signals. For more information see Algorithm 1,SI.

1. For an adequate duration, say 100 successive steps, let the dynamic of Eqs.3 and 4, upon which the amount of determinant agents evolve, proceed. Here, we reckon that the signaling patterns of *s*_1_ and *s*_2_ simultaneously evolve based on Eqs.1 and 2 and provoke the stem cells toward a possible destiny.
2. Follow up the “potential” destiny of the stem cell located at each grid on the plane. Allocate a 6*×* 6 look-up table for each of the present spices (just once at the very first iteration). We scale up the amounts of signals to the range of (0, 5). This is the variation interval of reverse spot signals. To allocate a likely fate to the cell, the signal concentrations are rounded up to the nearest integers, as the lookup table contains only discrete values. The rounded values of *s*_1_ are listed in the rows of each table, and the rounded values of *s*_2_ are listed in the columns. Then, Count the number of “virtual” offspring and renewed stem cells and categorize them based on the instant amount of *s*_1_ and *s*_2_ in each location, see Fig.3. Next, insert the statistic into the corresponding look-up table in each case. It is crucial to highlight that at this point the fate of the cells is not definitely determined. Instead, an assessment of their potential fate can be derived by taking into account the spatiotemporal value of (*s*_1_, *s*_2_).
3. Repeat the two previous steps for 1000 times and record the corresponding classified data according to the upper mentioned method. In this way, one collects more data and in consequence, the final predicted fate of the cells is closer to that of a real system.
4. Once every thousand steps, assess the amount of *s*_1_ and *s*_2_ on every single grid of the main substrate and find out the corresponding *s*_1_ and *s*_2_ on each of the six look-up tables. Then, compute the probability of the virtual emergence of each species simply as the number of that species in the look-up table, divided by the number of the whole population. After calculating the probability of all possible outcomes of the cell fate random variable, we compare them with the same quantities for the 1000 steps ahead and replace their maximum difference in *d* variable, which is taken as an arbitrarily large value that guarantees that we will have enough repetitions in our simulations. Repeat the above sequence of instructions until the amount of *d* is less than that of *e* = 0.0025, which is adopted as an arbitrary and constant limit for the acceptable error in our simulations.
5. Finally, substitute the initial distribution of mother cells *SC*_1_ and *SC*_2_ on the substrate with the final pattern of daughter cells of each type based on their come-up probability. At this stage, every single grid of the main substrate is implanted with the species that more likely conform with the influence of signal agents pair (*s*_1_, *s*_2_) on the internal switch of determinants, see Fig.3. For a summary of the ordered process see Algorithm 1,SI.

#### Algorithm 1: The sequential instruction to form a complex cellular pattern based on a given signaling blueprint

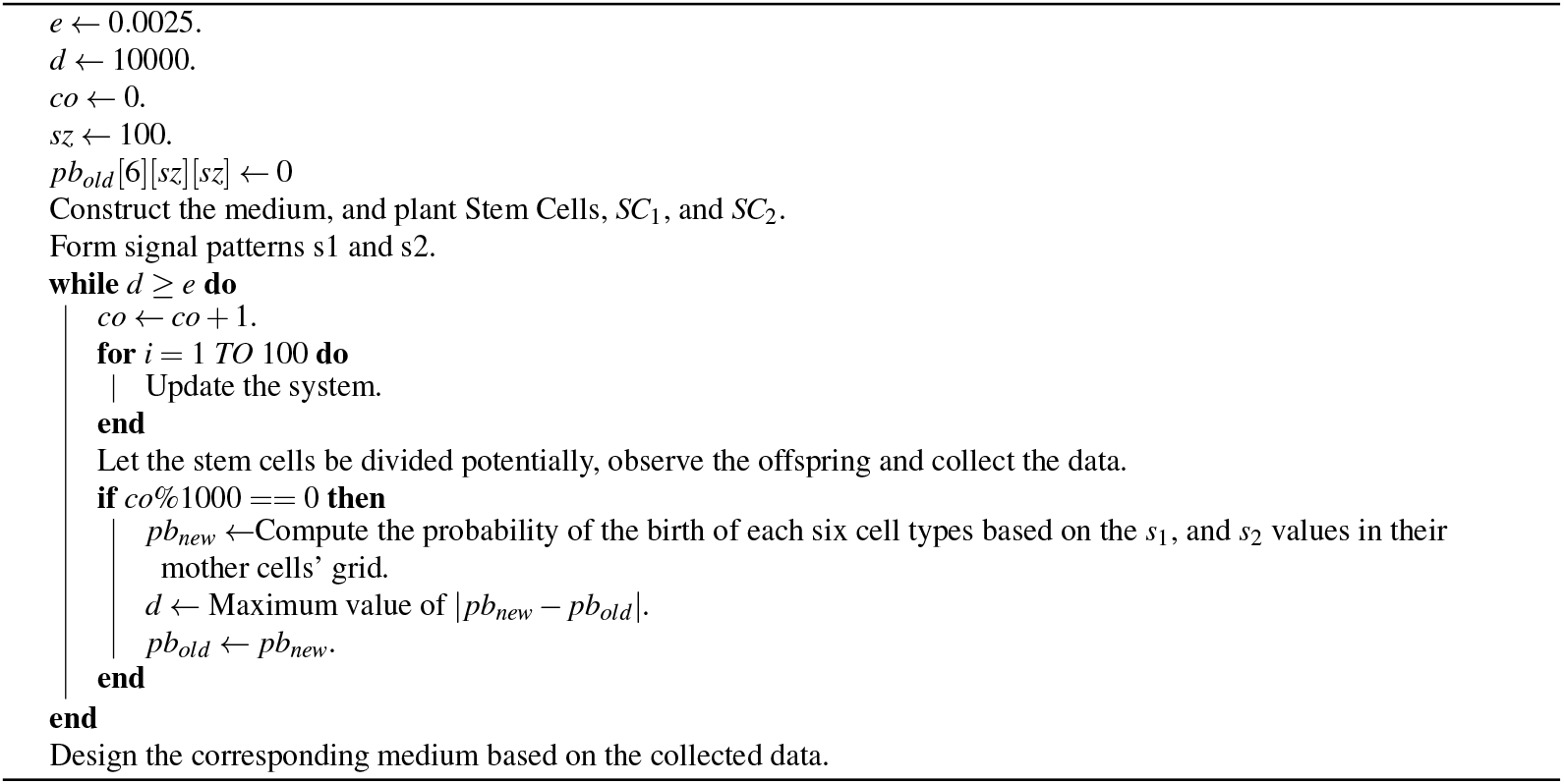

## Results

### Our signal-dependent tri-stable switch works

Fig.4(d) illustrates the solutions of Eq.3 and Eq.4 in the presence of variants pairs of (*s*_1_, *s*_2_). From different columns of the figure, it is evident that when the stem cells are exposed to different pairs of (*s*_1_, *s*_2_) the following fate of cell differentiation differs. The stem cells’ response to the presence of signals, which is implemented via Eq.5 and Eq.6, depends on the amount of both *s*_1_ and *s*_2_. In other words, there are pair combinations of (*s*_1_, *s*_2_) by which *SC*_1_ is become influenced, while *SC*_2_ remains neutral .*e*.*g*, column 1 to 4, and *vice versa* (.*e*.*g*, column 5 to 8). On the other hand, every single stem cell differentiates to one of its potential offspring based on the amount of (*s*_1_, *s*_2_) to which it has been exposed to. The first row of the tabular Fig.4(d) displays the final course of action of *SC*_1_ in the presence of variant combinations of (*s*_1_, *s*_2_). As it is seen in this row, in the presence of (*s*_1_, *s*_2_) = (0, 0) *B* species is superior. The same trend is seen when (*s*_1_, *s*_2_) = (1.5, 1.5) but with less difference between *A* and *B* production. In the presence of (*s*_1_, *s*_2_) = (2.5, 2.5) the process is reversed and *A* production becomes prior to that of *B* cells. When *SC*_1_ experiences (*s*_1_, *s*_2_) = (4, 4) pair signals, *A* species becomes superior. The corresponding signal pairs of the last four columns have no impact on precessing one of the species. Similarly, the second row illustrates the behavior of *SC*_2_ in the presence of different pairs of (*s*_1_, *s*_2_). It is seen that the first four rows have no specific influence on altering production probabilities of *C* and *D*. While in the presence of (*s*_1_, *s*_2_) = (0, 4) production rate of *C* is higher, the process becomes reversed when (*s*_1_, *s*_2_) pairs reaches to (2.5,1.5). When *s*_2_ vanishes and *s*_1_ is on its highest value,*e*.*i*. 4, production probability of *D*(*C*) is maximum (minimum).

**Figure 4.**
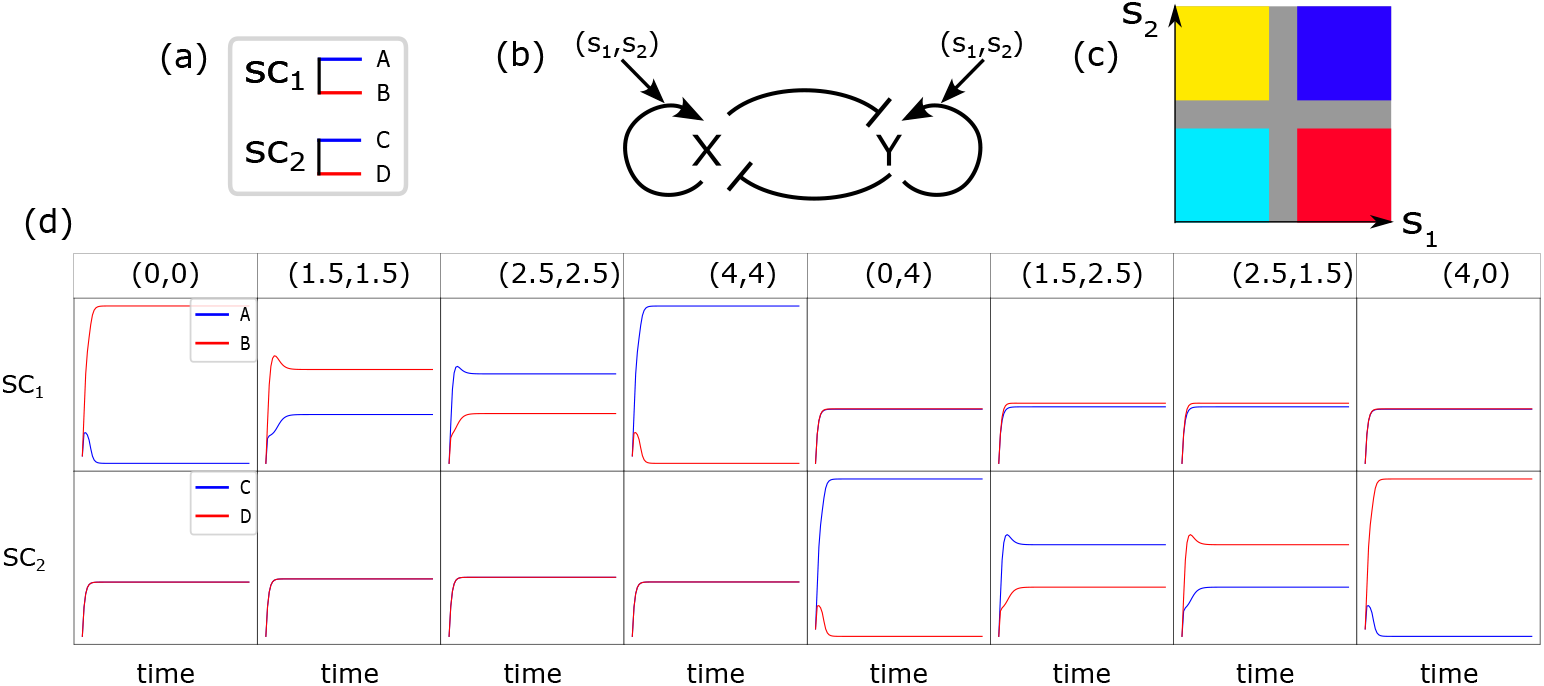
Division of *SC*1 result in either to *A* or *B* and the upshot of *SC*2 is either *C* or *D*, depending on the amount of signaling agent that each individual cell is subject to(a).The regulating switch inside each stem cell which contributes to developing it toward a special fate under the influence of environmental stimulant pairs of (*s*_1_, *s*_2_) (b). The cruciform shape of the phase field of possible developed cells in (*s*_1_, *s*_2_) plane. Blue and cyan colors render phenotypes *A* and *B* while yellow and red colors refer to *C* and *D* phenotypes, respectively. (c). The fate of each stem cell SCi (i = 1,2) is affected by exposing a special combinations of (*s*_1_, *s*_2_) pairs (d).

### Individual cellular decisions lead to collective cellular patterns under the influence of combined signals

Fig.5 depicts the arrangement of the final developed cellular patterns induced as a result of variant possible combinations of signaling patterns governed by Eq.1 and Eq.2. The color bar represents different cell types. Purple and green stand for *SC*_1_ and *SC*_2_, respectively. Blue and cyan colors render phenotypes *A* and *B* while yellow and red colors refer to *C* and *D* phenotypes, respectively. Each array of this arrangement corresponds to the combination of two members of the solution family Eq.1 and Eq.2, which are depicted inline in each case, see next paragraph. According to Fig.2, the solution family of these equations has three members: spot (left panel), reverse spot (middle panel), and stripe (right panel). From Fig.5, It is evident that the combination of these signaling patterns leads to a diverse collection of distinctive cellular patterns.

**Figure 5.**
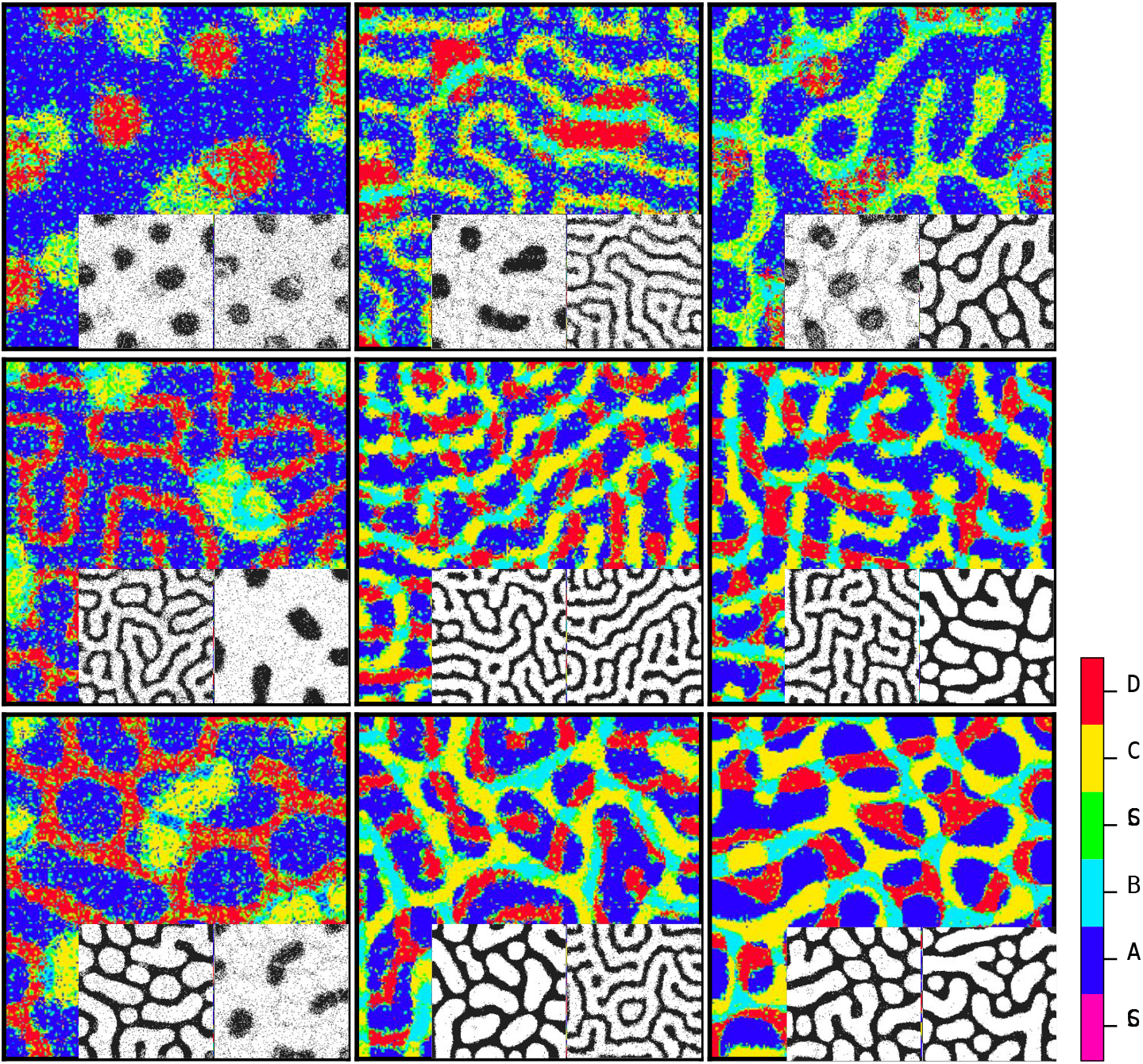
Steady state patterns of developmental cellular arrangements in a competitive signaling field. The patterns have been formed by the means of consecutive implementing of the algorithm 1.

### One can recognize the signal patterns from the final cellular arrangement, provided that the prior assumption of the pattern is given

Fig.5 illustrates the final cellular arrangements of different combinations of signaling patterns which are produced by Eq.1 and Eq.2. The corresponding acquired signaling patterns are depicted in each array, inline. Recognition of signal patterns is a directional process:

First, it is necessary to consider two blank planes, each of which in accord with one of the signals to project its corresponding pattern onto it. Next, we go through every single pixel of the cellular pattern. Then, based on the color of the pixel, we map the projection of this color onto the signal planes. Let us assume that the color of a pixel is blue, meaning that this pixel is occupied with a cell of phenotype *A*. According to the relations 5 and 6 as well as Fig.3, this implies that at this spot, the concentration of both signals is around its own summit. As a result, the projection of every blue pixel of the cellular pattern on both signal planes is a white point. Similarly, cyan color in the cellular pattern corresponds to *B* phenotype, whose occurrence is highly probable when the concentration of both signals is low. Accordingly, the map of each cyan pixel matches a corresponding black color on both signal planes. Likewise, the yellow (red) color represents *C* phenotype (*D* phenotype), whose production rate is high when the concentration of *s*_1_ is low (high), while that of *s*_2_ is high (low). As a consequence, the projection of each yellow pixel onto the corresponding point on the first signal plane is white (black) while its projection onto the similar point on the second signal plane is black (white). For a summary of the ordered process see Algorithm 2, SI.

## Discussion

The positional stimuli have been emerging as key regulators of transcription and gene expression in diverse physiological contexts^26^. These environmental drivers engage in the phenotypic diversity and proliferation/differentiation balance of stem cells^23,27,28^. The regulation process of non-genetic diversity involves the interplay of intracellular as well as intercellular components to interpret positional cues^29,30^. In a competing arena in which various chemical stimulants vie for affecting more on cell’s fate, the process demands more robust and complex mechanisms. In order to specify and extend their offspring territory, the stem cells utilize a signaling process to communicate and collaborate with each other. This process ends in collective self-organized forms on length scales that are much larger than those of the individual units^31^.

In this study, first, we investigated the impact of passive competing external signals on intracellular switches of a single stem cell. This provides us with a direct inspection of the connection between intracellular and extracellular dynamics. Indeed, by mapping the environmental signaling patterns to the probability of the emergence of differentiated cell types, this model is capable of capturing any desired complex pattern, whether passive or active. The sort of models that recapitulates signaling dynamics and predicts cell fate patterning upon chemical perturbations preceedingly has been investigated in non-competitive environments^15,25,32–36^. However, here, we focused on the behavior of each cell in interaction with diverse and competing signals. Fig.1 illustrates the resulting phenotypic cellular patterns of different combinations of two typical signal profiles of Gaussian and sinusoidal blueprints.

The environmental signals influence the fate of each stem cell, SCi (i = 1,2), by means of biasing the regulation of our tri-stable switch, see Eqs.3 and 4. Based on the definition of 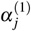 and 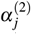 Eqs.5 and 6, it is evident that the pairs of (*s*_1_, *s*_2_) are relevant in controlling decisions of this switch. This definition is advantageous in variant aspects: First, it directs each stem cell’s fate to the symmetric phase space of Fig.4(c), in which each of the patches corresponds to one of the resulting phenotypes and there is no dominant domain between them. Besides, the representative patches are far enough apart that lead to distinctive outcomes in the occurring cellular pattern field. The narrow cruciform band, *i*.*e*, the grey area in Fig.4(c), between these four patches is

### Algorithm 2: The sequential instruction to speculate about the triggering signalling patterns corresponding to the cellular arrangements

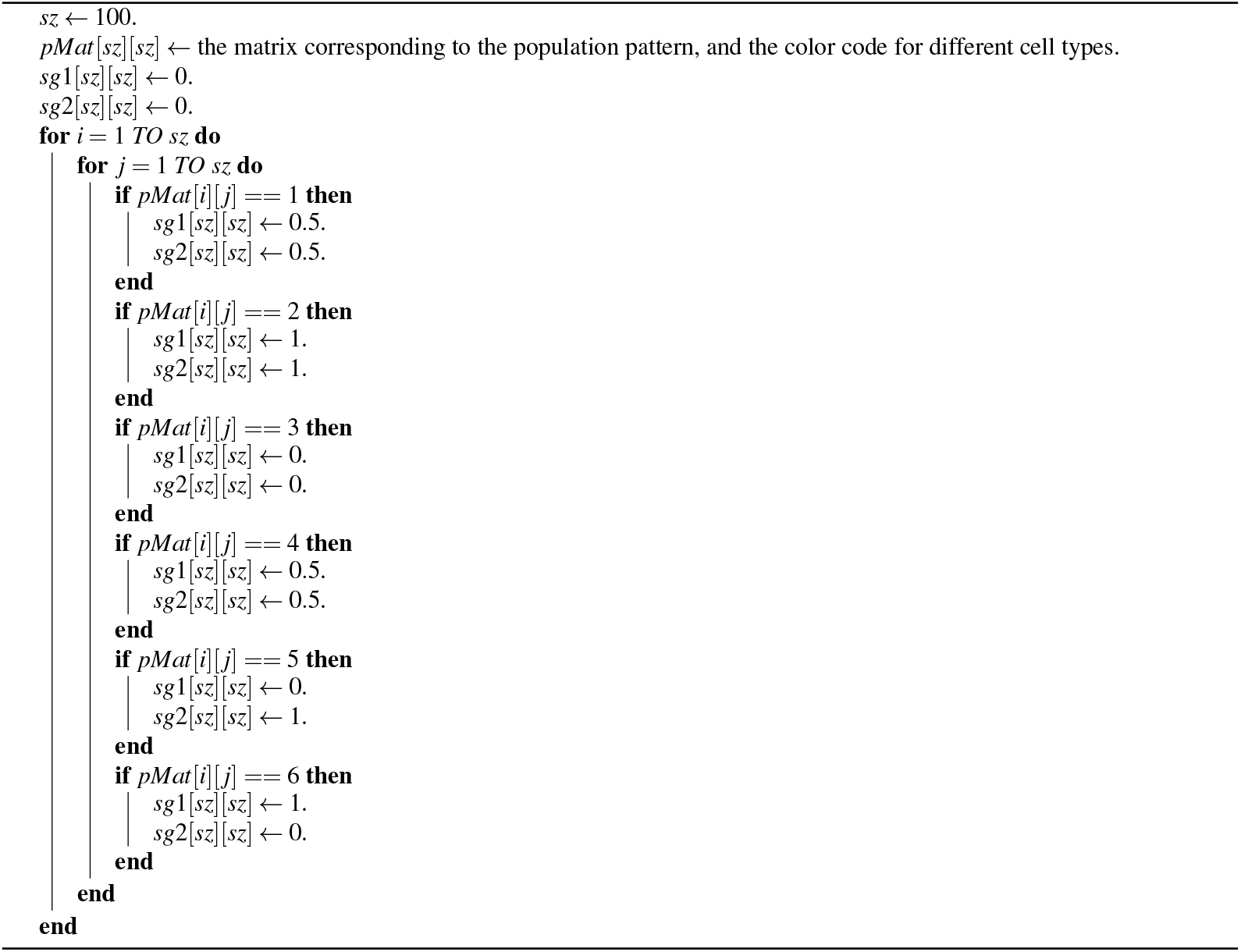

where the fate of each cell is determined stochastically. From Fig.4(d), it is evident that the regulatory switch plays either an active role or a neutral one based on the amount of existing signals (*s*_1_, *s*_2_) in each point, *i*.*e*, combinations of (*s*_1_, *s*_2_) which effectively lead to offspring *A* or *B* from *SC*_1_ have nothing to do with *SC*_2_ and *vice versa*. In consequence, there is a smooth transition from left to right in each row of Fig.4(d).

After investigating the impact of static environmental stimulants on the internal switch, we dealt with the active signaling between the sources of producing variant phenotypes. We took advantage of confined Turing models for two different signals secreted from each of stem cells^21^. The dynamic includes linear reaction terms and additional constraints that confine the two variables within a finite range. The resulting patterns of this dynamic are either stationary striped patterns or spotted patterns. The second pattern, in turn, consists of two forms: spotted and reverse spotted patterns. Here, the tuning parameter upon which the pattern type is specified is the maximum concentration of the activator 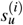 *i ∈*1, 2^21^, see Fig.2. Based on this prior dynamic, nine distinct mutual patterns are generated by the two signals 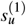 and 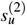 Stochasticity has been proven to be another non-genetic diversifying resource of variation in nature^30,37–41^. It has shown that controlled amount of randomness ends in phenotypic variation and as a result, population heterogeneity^25,42,43^. In this study, to reflect the non-deterministic portion of the signaling system, we implemented the Gillespie algorithm^22^ by stepping in time to successive molecular reaction events according to the premises of the Shoji, *et*.*al*, model^21^, see Eqs.1 and 2. Another aspect of incorporating randomness in our reductionist insight is simulating the emergence of every species in the look-up table based on the calculation of its corresponding probability, see Fig.3.

We see that the synthesis of the signaling arrangement with the switch in the presence of controlled noise creates a rich highly reproducible organizations of differentiated cells. Fig.5 depicts the resulting patterns of the differentiated cell which have been exposed to various combinations of active signaling lay-outs of Fig.2. The procedure we dealt with in this study is one of the various known roots to construct an organized arrangement of cells. Mobility of cells^44^, modulation of the physical and geometrical environment^16^, and priming with chemical signals^1^ are among other intrinsic capacities of stem cells to make patterns. In practice, a combination of all these methods incorporates to form an organization. Nevertheless, it is seen that solely following chemical environmental cues leads to the production of a rich and wide range of patterns.

The systematic approach that we proposed in this study, enables us to simulate the final configuration of a complex organization in terms of basic biophysical processes. The algorithm we followed here, leaps over many details, *e*.*g*, the dynamic environment in the presence of actual division, born of different types of cells, apoptosis, cell movement, *etc*. Nonetheless, the final structure of the pattern of differentiated cells in a competitive environment that is offered by the method is statistically reproducible. Moreover, given that the final state of the cellular organization is known, the corresponding signaling patterns based on which the cellular organization constructed is partly predictable following the provided process. Importantly, for any given complex cellular pattern for which merely the prior class of signal patterns is known, the provided method concludes closely the signaling profile that set off the cellular pattern.

## Author contribution

M.S. designed research; N.K. and Z.E. performed research and analyzed data; Z.E. wrote the original version of the manuscript; Z.E., N.K., and M.S. contributed to manuscript revision.

## Additional information

The authors declare no competing interests.

